# Dual α-globin and truncated EPO receptor knockin restores hemoglobin production in α-thalassemia-derived red blood cells

**DOI:** 10.1101/2023.09.01.555926

**Authors:** Simon N. Chu, Eric Soupene, Beeke Wienert, Han Yin, Devesh Sharma, Travis McCreary, Kun Jia, Shota Homma, Jessica P. Hampton, James M. Gardner, Bruce R. Conklin, Tippi C. MacKenzie, Matthew H. Porteus, M. Kyle Cromer

## Abstract

Alpha-thalassemia is an autosomal recessive disease with increasing worldwide prevalence. The molecular basis is due to mutation or deletion of one or more duplicated α-globin genes, and disease severity is directly related to the number of allelic copies compromised. The most severe form, α-thalassemia major (αTM), results from loss of all four copies of α-globin and has historically resulted in fatality *in utero*. However, *in utero* transfusions now enable survival to birth. Postnatally, patients face challenges similar to β-thalassemia, including severe anemia and erythrotoxicity due to imbalance of β-globin and α-globin chains. While curative, hematopoietic stem cell transplantation (HSCT) is limited by donor availability and potential transplant-related complications. Despite progress in genome editing treatments for β-thalassemia, there is no analogous curative option for patients suffering from α-thalassemia. To address this, we designed a novel Cas9/AAV6-mediated genome editing strategy that integrates a functional α-globin gene into the β-globin locus in αTM patient-derived hematopoietic stem and progenitor cells (HSPCs). Incorporation of a truncated erythropoietin receptor transgene into the α-globin integration cassette dramatically increased erythropoietic output from edited HSPCs and led to the most robust production of α-globin, and consequently normal hemoglobin. By directing edited HSPCs toward increased production of clinically relevant RBCs instead of other divergent cell types, this approach has the potential to mitigate the limitations of traditional HSCT for the hemoglobinopathies, including low genome editing and low engraftment rates. These findings support development of a definitive *ex vivo* autologous genome editing strategy that may be curative for α-thalassemia.

**Graphical abstract:** 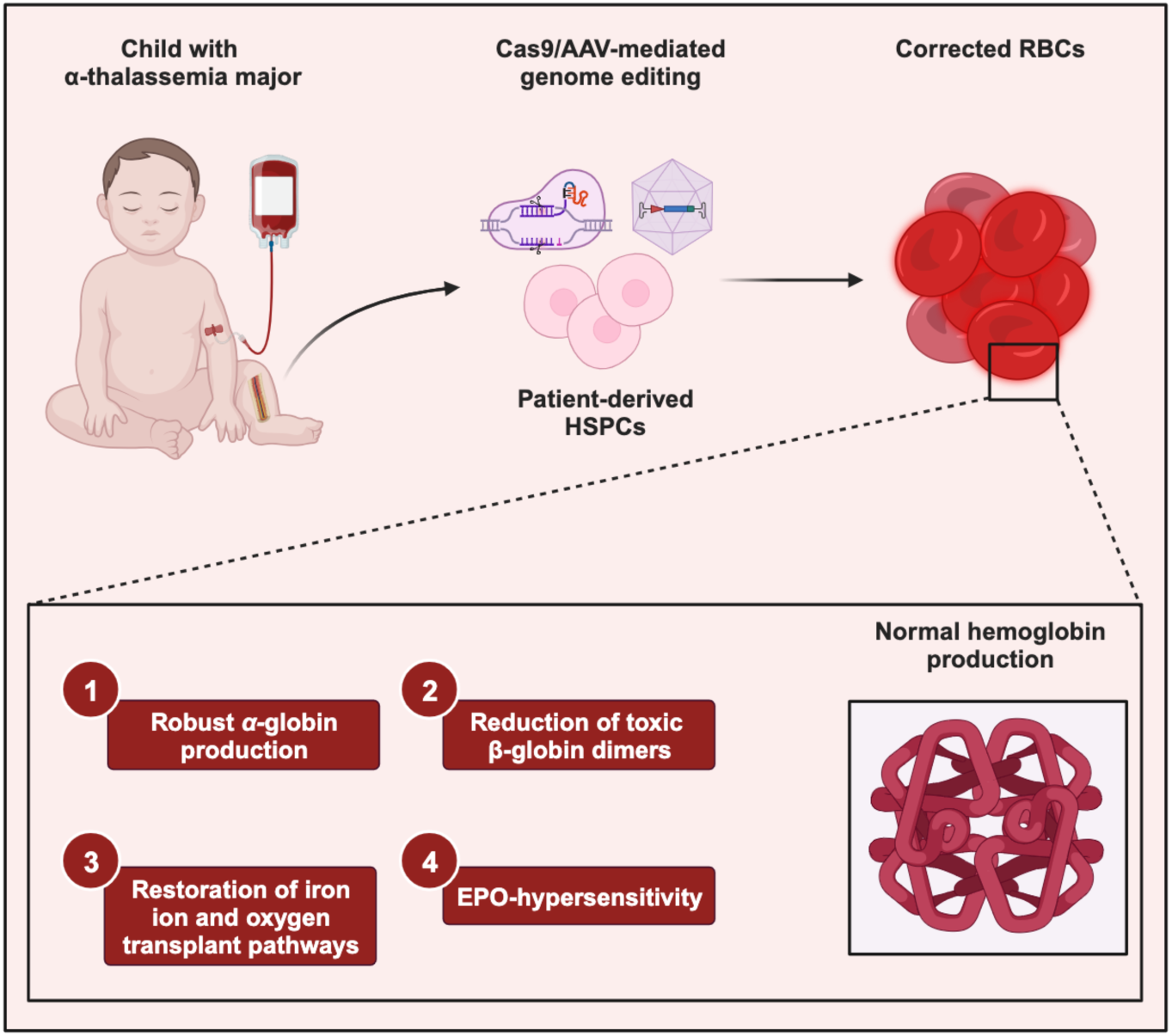

## Introduction

Alpha-thalassemia is one of the most common monogenic diseases in the world. It is currently estimated that 5% of the world’s population harbors an α-thalassemia variant^1^. While the carrier frequency is highest in those with Southeast Asian heritage, data suggests that these rates are rising due to population growth, human migration, and advances in treating milder forms of the disease^2^. Humans harbor four copies of α-globin genes (a direct tandem repeat on chromosome 16), and disease severity directly correlates with the number of mutated or deleted alleles. The most severe form of the disease, in which all four copies are disrupted, is called α-thalassemia major (αTM). While this disease has historically been lethal *in utero*, patients may now survive to birth after *in utero* blood transfusions, often with excellent neurologic outcomes^3,4^. Postnatally, these patients present with a disease similar to β-thalassemia, with the absence of α-globin leading to an inability to form hemoglobin heterotetramers and subsequent severe anemia. Moreover, the accumulation of orphan β-globin chains, which form oxidized, covalently linked dimers and an unstable homotetramer (Hemoglobin H), leads to erythrotoxicity and hemolysis^5^. As a consequence, these patients require chronic transfusions, which often results in iron overload and need for iron chelation therapy^6^. Although allogeneic-hematopoietic stem cell transplantation (HSCT) may provide a cure, suitable matched donors are only available in a minority of cases and carry a risk of immune rejection and graft-versus-host-disease (GvHD)^7^. Furthermore, while numerous gene therapy and genome editing strategies have been developed for patients with β-thalassemia^8,9,10^ there are no such therapies for the most severe forms of α-thalassemia, indicating a major unmet medical need for this patient population.

In this work, we developed a novel Cas9/AAV6-mediated genome editing strategy to integrate a functional copy of the *α-globin* gene into the *β-globin* locus in αTM patient-derived hematopoietic stem and progenitor cells (HSPCs) as a universal treatment strategy for the disorder—an approach that could be curative for any αTM patient regardless of the specific causative mutations or deletions. This approach allows both correction of the underlying disease as well as restoration of the β-globin:α-globin imbalance by placing the *α-globin* transgene under erythroid-specific expression of the *β-globin* locus. Taking a cue from human genetics, we also incorporated a naturally occurring truncated erythropoietin receptor (*tEPOR*) cDNA into the *α-globin* integration cassette, allowing for simultaneous correction of α-thalassemia and increased erythropoietic output from edited HSPCs^12^. By directing edited HSPCs toward increased production of clinically relevant RBCs instead of other divergent cell types, this approach has the potential to overcome many of the clinical challenges of HSCT for treatment of the hemoglobinopathies, including low editing and engraftment rates, as well as high morbidity from prerequisite myeloablative regimens^13,14,15^. Similar to our prior work with β-thalassemia^8^, this strategy could be used in to develop an autologous-HSCT treatment for αTM, overcoming the shortage of matched donors and ameliorating the risk of immune rejection and GvHD for patients suffering from this disease.

## Results

### Efficient cleavage with HBB intron-targeting gRNAs

Due to the high editing frequencies achieved in primary HSPCs in prior work^8,16^, we sought to use a Cas9/AAV6-mediated genome editing strategy to knock an *α-globin* (*HBA*) transgene into the *β-globin* (*HBB*) locus in patient-derived HSPCs. To do so, we first designed and screened Cas9 gRNAs at the *HBB* locus. So that cleavage alone without homology-directed repair (HDR) would not disrupt β-globin production, we chose 22 candidate gRNAs located in intron 1 or intron 2 as well as the 3’ UTR (Figure 1A and Supplemental Table 1). To determine cleavage frequencies, we delivered candidate gRNAs pre-complexed with Cas9 protein to the human HUDEP-2 cell line via electroporation^17^. Genomic DNA was harvested several days post-targeting and subjected to PCR amplification of the region surrounding the expected cleavage site. Insertion and deletion (indel) frequencies of the corresponding Sanger sequences were then quantified using ICE analysis^18^. We found that targeting the 5’ and 3’ UTR regions of *HBB* proved not to be feasible, as the former had significant homology to δ-globin and the latter had no gRNAs with detectable cleavage. We found that the most efficient gRNAs corresponded to sg7 in intron 1, and to sg11 and 13 in intron 2 (Figure 1A and Supplemental Figure 1)^19^. *In silico* off-target analysis of these guides using COSMID^20,21^ revealed that sg7 had the most favorable predicted off-target profile (Figure 1B). On further evaluation of the 19 predicted off-target sites for sg7, all were found to reside in non-coding regions of the genome (Figure 1C).

**Figure 1.**
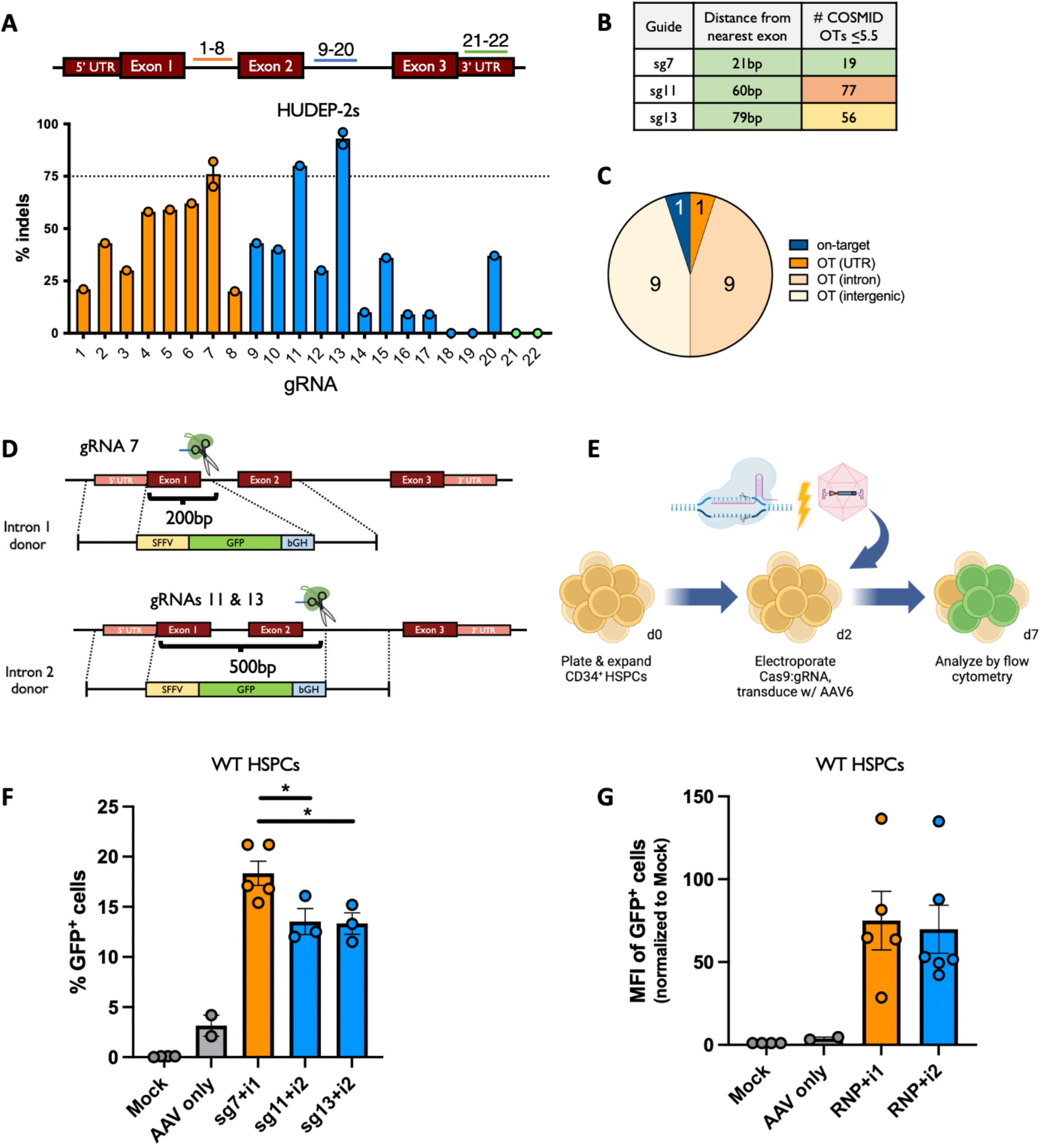
Efficient integration of HDR templates at *HBB* locus. (A) Schematic of *HBB*-targeting gRNAs and resulting indel frequencies in HUDEP-2 cells following electroporation-mediated delivery of Cas9:gRNA RNPs. Dotted line indicates indel frequencies at 75% threshold. Bars represent mean ± standard error of the mean (SEM). (B) *In silico* off-target analysis of gRNA 7, 11, 13 using COSMID. (C) Pie chart displaying genomic features at 19 predicted target sites for sg7. (D) Schematic of custom AAV6 DNA repair donors designed to mediate integration at intron 1 using sg7 or intron 2 using sg11 or 13. Bracket indicates distance between left and right homology arms. (E) Schematic of Cas9/AAV6 genome editing workflow in primary HSPCs. (F) Percentage of GFP^+^ cells following editing in WT HSPCs was determined at d5 post-editing using flow cytometry. Bars represent mean ± SEM. * denotes *p*<0.05 by unpaired two-tailed t-test. (G) Mean fluorescence intensity (MFI) of GFP^+^ cells from Figure 1F was determined by flow cytometry. Bars represent mean ± SEM.

### Efficient integration of HDR templates at HBB locus

Following identification of effective *HBB* intron-targeting gRNAs, we developed DNA repair templates packaged in AAV6 delivery vectors that could effectively mediate HDR at these cleavage sites (termed Intron 1 and Intron 2 donors; Figure 1D). Each integration cassette was comprised of a SFFV promoter driving expression of a GFP reporter to allow rapid readout of integration frequencies via flow cytometry. While the right homology arm of each DNA repair template corresponded to the ∼900bp immediately downstream of the intron 1 or intron 2 Cas9 cut site, the left homology arm was split away from the cleavage sites to correspond to the ∼900bp immediately upstream of the start codon of the endogenous *HBB* gene. As in a prior study^8^, this split homology arm strategy is expected to allow promoterless integration cassettes—ultimately an *α-globin* transgene in this work—to be driven by the regulatory machinery of the endogenous locus.

Once DNA repair templates were assembled and packaged into AAV6 vectors, we tested each integration strategy by delivering the most effective gRNAs (sg7, 11, or 13) complexed with high-fidelity Cas9 protein^22^ to wild-type (WT) human primary CD34^+^-enriched HSPCs via electroporation. immediately following electroporation, we transduced cells with AAV6 vectors corresponding to either intron 1 or intron 2 integration schemes. Five days later, we analyzed targeting frequencies by flow cytometry (Figure 1E and Supplemental Figure 2A) and found that the Intron 1 integration strategy was significantly more efficient (median of 17.1% GFP^+^ cells) compared to Intron 2 integration (median of 12.5% for sg11 and 13.3% for sg13; *p*<0.05) (Figure 1F), perhaps due to the shorter distance that the left homology arm was split away from the Cas9 cleavage site. In addition, both integration strategies yielded a high mean fluorescence intensity (MFI) per edited cell (Figure 1G). Given the high cleavage and integration frequency, as well as the favorable off-target profile of sg7, we proceeded with further testing in primary WT HSPCs using the Intron 1 integration strategy.

### Integration at HBB drives erythroid-specific expression of transgenes

A key goal of this work is to achieve erythroid-specific expression of an *α-globin* transgene. To test whether the *HBB* Intron 1 integration strategy is able to achieve this, we designed custom promoterless integration cassettes which included either the full-length *HBA1* or *HBA2* transgene (i.e., including all UTRs, exons, and introns) linked to a T2A-YFP reporter to allow quantification of HDR frequency by flow cytometry and to serve as a surrogate for α-globin protein production (Figure 2A). Because the specific UTRs flanking an integrated transgene can have a great bearing on expression^8^, we also designed integration strategies that would allow our *HBA* transgene to be flanked by *HBB*, *HBA1*, and *HBA2* 5’ and 3’ UTRs.

**Figure 2.**
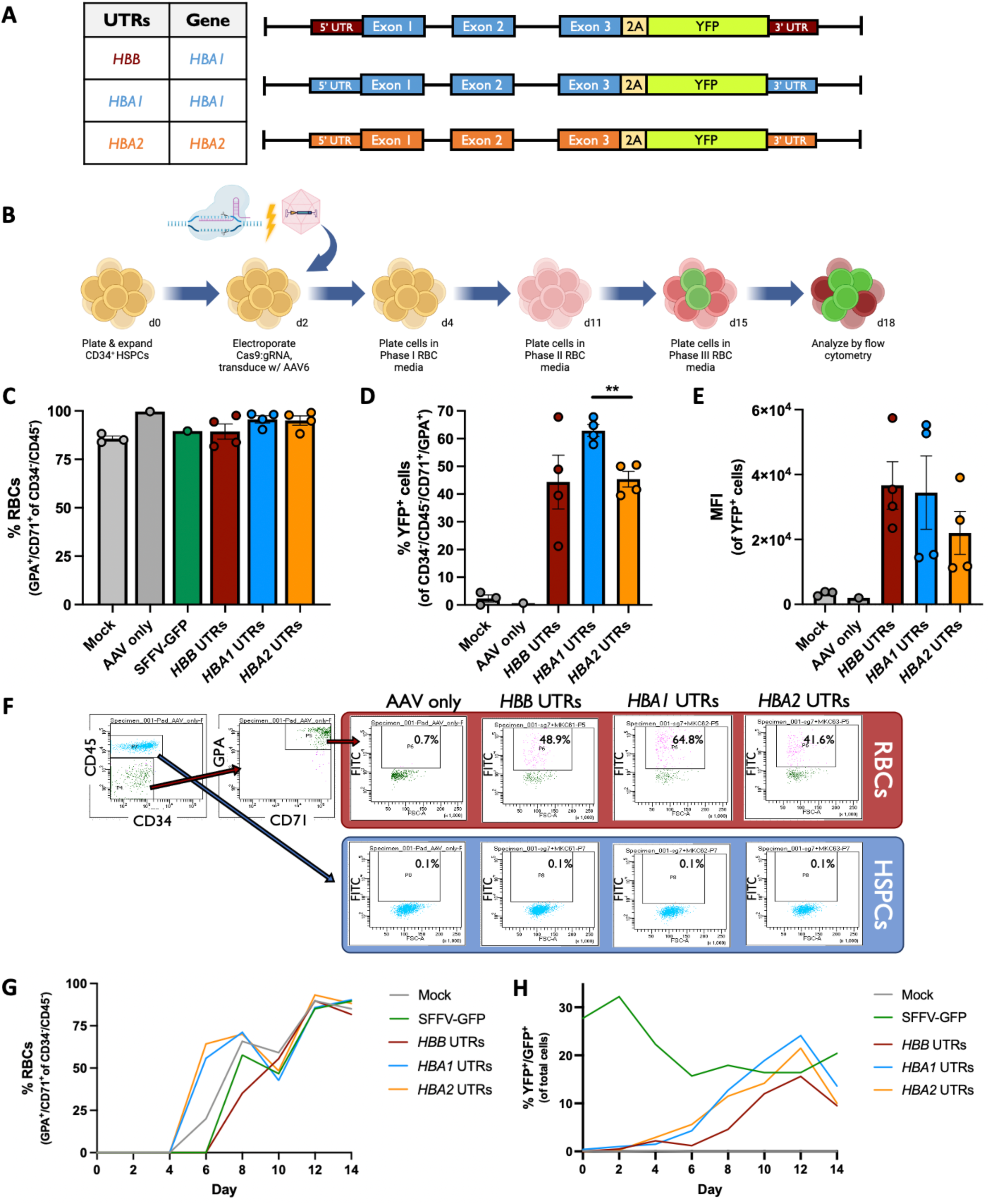
Integration at *HBB* drives erythroid-specific expression of transgenes. (A) Schematic of custom AAV6 DNA repair donors designed to integrate promoterless *HBA-2A-YFP* transgenes at the start codon of *HBB*. Table to the left indicates whether cassette integrated *HBA1* or *HBA2* transgene as well as *HBB*, *HBA1*, or *HBA2* UTRs. (B) Schematic of Cas9/AAV6 genome editing workflow in primary HSPCs followed by *in vitro* RBC differentiation. (C) Percentage of CD34^−^/CD45^−^ HSPCs acquiring RBC surface markers—GPA and CD71—as determined by flow cytometry. Bars represent mean ± SEM. (D) Percentage of YFP^+^ cells in CD34^-^/CD45^-^/CD71^+^/GPA^+^ cells was determined at d14 of RBC differentiation using flow cytometry. Bars represent mean ± SEM. ** denotes *p*<0.005. (E) MFI of YFP^+^ cells from Figure 2D was determined by flow cytometry. Bars represent mean ± SEM. (F) At d11 of RBC differentiation, cells were stained for HSPC/RBC markers and analyzed by flow cytometry. Percentage of YFP^+^ cells are noted (N=1) (G) Percentage of CD34^−^/CD45^−^ cells that acquired CD71 and GPA RBC markers are plotted over the course of RBC differentiation (N=1). (H) Percentage of YFP^+^ or GFP^+^ cells are plotted over the course of RBC differentiation (N=1).

To determine whether integration of these promoterless *α-globin* cassettes at *HBB* yielded erythroid-specific expression, we edited WT HSPCs using electroporation-mediated delivery of Cas9:sg7 RNP followed by transduction with AAV6 vectors containing each repair template. We then performed an established, 14-day erythroid differentiation protocol^23^ and quantified YFP expression and RBC differentiation (staining with antibodies for CD34, CD45, CD71, and GPA) via flow cytometry (Figure 2B and Supplemental Figure 2B). We found that all conditions efficiently differentiated into erythroid cells (Figure 2C). However, we did observe differences among the integration vectors in terms of HDR frequency and MFI of edited cells. We found that the vector integrating the *HBA1* transgene and *HBA1* UTRs achieved the most highly HDR-edited population of RBCs (median of 63% YFP^+^ cells; Figure 2D) and the second highest MFI of the HDR-edited cell population (Figure 2E). As confirmation that these integration strategies achieved erythroid-specific transgene expression, we observed high YFP fluorescence in cells that acquired RBC markers (CD71^+^/GPA^+^) in all editing conditions but no fluorescence above background in cells retaining HSPC markers (CD34^+^/CD45^+^) (Figure 2F). We also tracked differentiation and YFP expression kinetics and found that all editing conditions proceed through RBC differentiation at a similar rate (Figure 2G). When monitoring expression of fluorescent reporters, we find that cells edited with the constitutive SFFV-GFP integration vector maintain high expression over the course of RBC differentiation, whereas all three conditions edited with HBA-2A-YFP constructs display increasing fluorescence that dovetails with RBC differentiation (Figure 2H). Collectively, these results indicate that integration of the *HBA* transgene at *HBB* achieves high levels of erythroid-specific transgene expression.

### α-globin integration in αTM HSPCs yields low levels of hemoglobin

While T2A-YFP reporters allow rapid readout of HDR-editing frequency, to generate clinically relevant constructs we created *HBA1* integration vectors without T2A-YFP that were flanked by either *HBB* or *HBA1* UTRs (Figure 3A). To test these new vectors, we edited WT HSPCs as before and found that both vectors mediate high frequencies of HDR editing (median of 36.6% and 42.3% HDR-edited alleles for *HBB* and *HBA1* UTR vectors, respectively) as measured by a custom ddPCR assay (Figure 3B).

**Figure 3.**
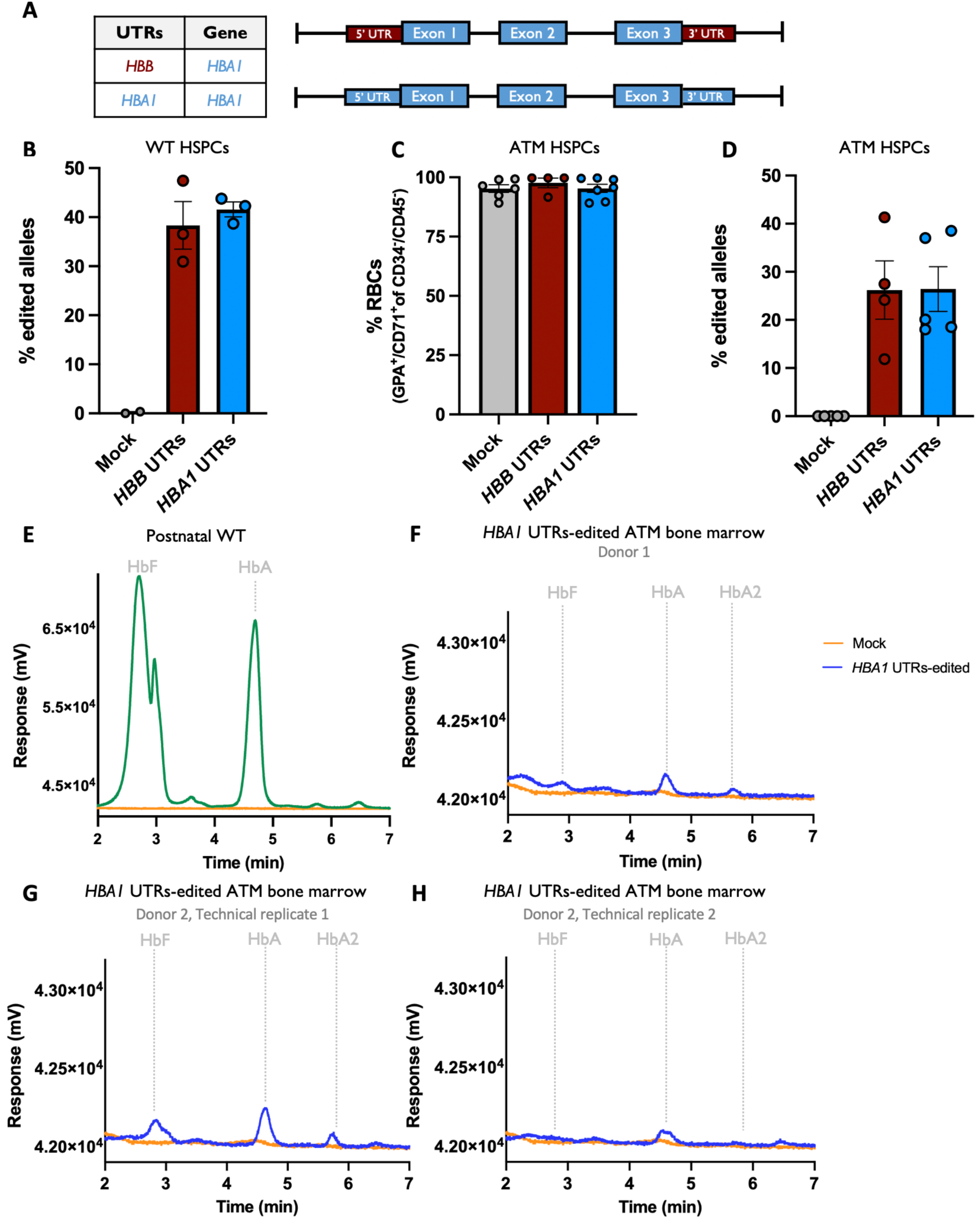
α-globin integration yields low levels of hemoglobin in αTM HSPCs. (A) Schematic of custom AAV6 DNA repair donors designed to integrate promoterless *HBA* transgenes at the start codon of *HBB*. Table to the left indicates whether the cassette integrated *HBA1* transgene with *HBB* or *HBA1* UTRs. (B) Percentage of edited alleles in WT HSPCs at d14 of RBC differentiation. Bars represent mean ± SEM. (C) Percentage of CD34^−^/CD45^−^ αTM HSPCs acquiring RBC surface markers—GPA and CD71—as determined by flow cytometry. Bars represent mean ± SEM. (D) Percentage of edited alleles in αTM HSPCs at d14 of RBC differentiation. Bars represent mean ± SEM. (E) HPLC elution chromatogram displaying the hemoglobin tetramer profile from WT healthy control HSPCs following *in vitro* RBC differentiation. Time displayed on the x-axis represents retention time in minutes for each hemoglobin tetramer type to elute. Absorbance on the y-axis indicates the concentration of a particular hemoglobin tetramer. (F-H) HPLC elution chromatograms displaying the hemoglobin tetramer profile from αTM HSPCs that have undergone editing and RBC differentiation. Chromatograms represent 2 different donors as well as a technical replicate from donor 2 that was edited independently.

We next sought to determine whether *α-globin* integration can restore hemoglobin tetramer production in patient-derived HSPCs following erythroid differentiation. We isolated CD34^+^ HSPCs from a bone marrow aspirate that was taken from a ∼1-year-old patient with αTM. We then edited and differentiated HSPCs as before and found that all conditions were able to effectively differentiate into RBCs (Figure 3C). Following ddPCR analysis, we found that cells were edited at a median of 25.8% and 20.1% HDR-edited alleles for *HBB* and *HBA1* UTR vectors, respectively (Figure 3D). We then performed hemoglobin tetramer HPLC analysis on edited patient-derived αTM cells at the end of erythroid differentiation and found that HDR-edited cells displayed a modest increase in all three alpha-containing hemoglobins (fetal hemoglobin (HbF), adult hemoglobin (HbA), and hemoglobin A2 (HbA2)) compared to WT (Figure 3E-H and Supplemental Figure 3). While these results demonstrate for the first time that gene therapy or genome editing may be used to increase α-globin production in αTM patient-derived RBCs, it is unlikely that this low level of α-globin production will yield significant clinical benefit.

### Dual integration of α-globin and tEPOR increases hemoglobin production in αTM HSPCs

Given that *α-globin* integration into *HBB* does not substantially restore hemoglobin production, we sought alternative strategies to increase the therapeutic potential of this approach. As described in a cohort that included an Olympic gold medalist cross-country skier^24^, a premature stop codon in the erythropoietin receptor (*EPOR*) was linked with a condition called congenital erythrocytosis, which is characterized by non-pathogenic hyper-production of RBCs. In subsequent work^12^, we demonstrated that genome editing-mediated integration of a truncated *EPOR* (*tEPOR)*-coding cDNA could increase erythropoietic output from primary human HSPCs. Thus, we hypothesized that co-expression of *tEPOR* cDNA from our *α-globin* integration cassette could lend a selective advantage to genome-edited and, therefore, α-globin-expressing erythroid cells. To test this, we added a *tEPOR* cDNA at the C-terminus of the *α-globin* integration cassette with expression linked by an internal ribosome entry site (Figure 4A). We then edited WT HSPCs with our *HBA1*-UTRs *α-globin* integration vector from Figure 3 and compared it against the bicistronic *α-globin*+*tEPOR* vector in WT HSPCs. In both instances, edited cells efficiently differentiated into erythroid cells (Figure 4B). While cells edited with *α-globin* integration alone maintained a constant editing rate (median of 39.2% edited alleles at d0 and 38.4% at d14), HDR frequencies increased significantly over the course of erythroid differentiation in cells edited with the *α-globin*+*tEPOR* cassette, rising from a median of 29.3% edited alleles at d0 to 69.9% at d14 (*** *p*=0.0004) (Figure 4C). As in prior work, this suggest that cells expressing a *tEPOR* have a significant competitive advantage over unedited cells during RBC differentiation. We next tested these vectors in αTM patient-derived HSPCs and again observed that edited and unedited cells were able to effectively undergo erythroid differentiation (Figure 4D). As with WT cells, we found that HDR-editing frequencies among cells targeted with the *α-globin* alone vector maintained a consistent editing rate over the course of RBC differentiation (Figure 4E). However, we again observed a dramatic increase in HDR frequencies in cells edited with the *α-globin*+*tEPOR* cassette, rising from a median of 40.5% HDR-edited alleles at d4 to 60.5% at d16 (*\* p*=0.02). In addition to the increase in editing frequencies, we also observed a 2-fold increase in cell counts at the end of RBC differentiation in conditions edited with the *α-globin*+*tEPOR* cassette compared to cells edited with the *α-globin* vector (** *p*=0.007) (Figure 4F), indicating that *tEPOR* expression is driving increased erythropoietic output from genome-edited HSPCs.

**Figure 4.**
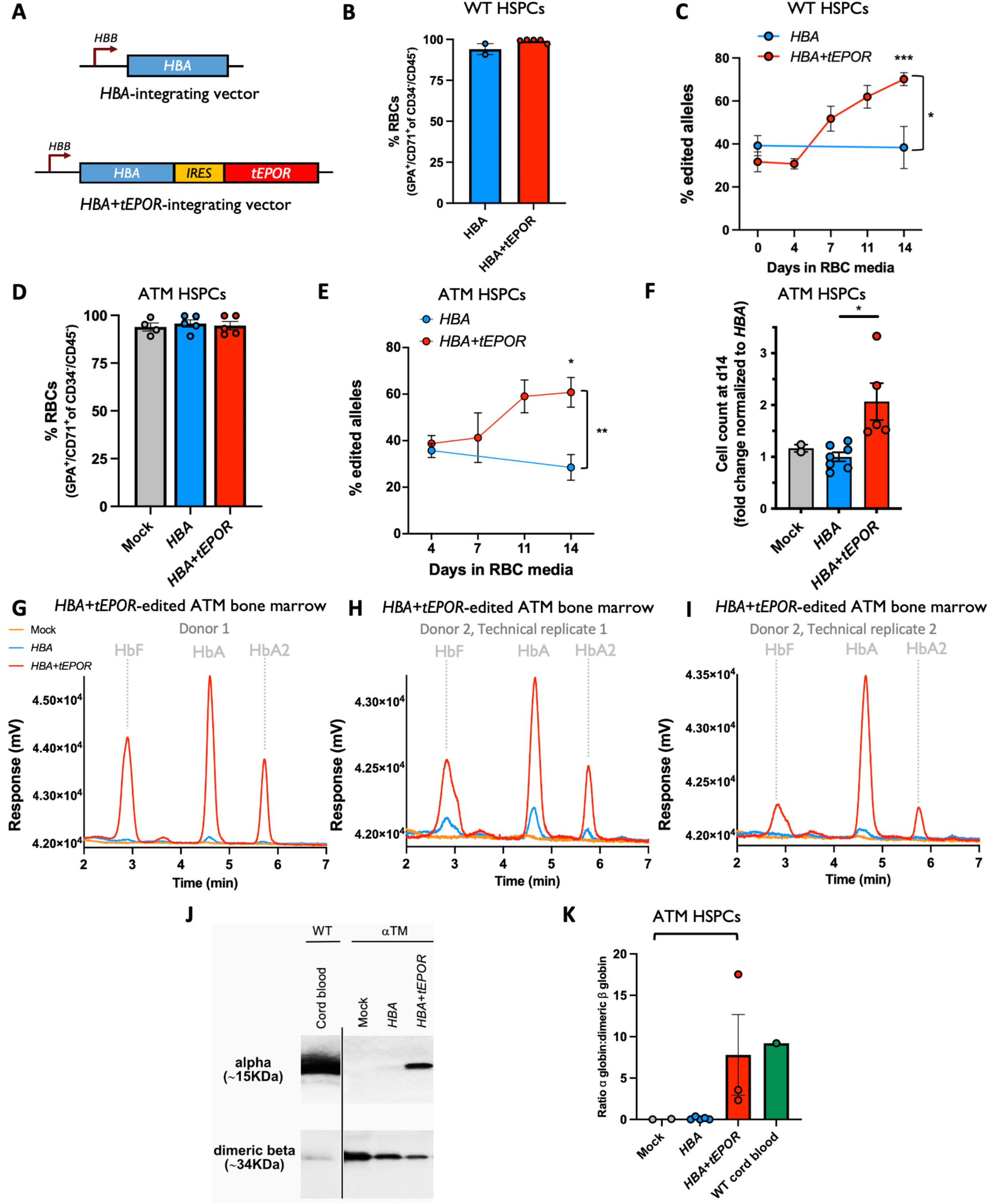
Dual integration of α-globin and *tEPOR* increases hemoglobin production in αTM HSPCs. (A) Schematic of custom AAV6 donors designed to integrate promoterless *HBA* and *HBA*+*tEPOR* transgenes at start codon of HBB. Both vectors are flanked by *HBA1* UTRs. (B) Percentage of CD34^−^/CD45^−^ WT HSPCs acquiring RBC surface markers—GPA and CD71—as determined by flow cytometry. Bars represent mean ± SEM. (C) Percentage of edited alleles in WT HSPCs over the course of RBC differentiation. Bars represent mean ± SEM. *** denotes *p*=0.0004 and * denotes *p*=0.01. (D) Percentage of CD34^−^/CD45^−^ αTM HSPCs acquiring GPA and CD71 as determined by flow cytometry. Bars represent mean ± SEM. (E) Percentage of edited alleles in αTM HSPCs over the course of RBC differentiation. Bars represent mean ± SEM. ** denotes *p*=0.009 and *\** denotes *p*=0.02. (F) Cell count at d14 of RBC differentiation with fold change normalized to *HBA*. Bars represent mean ± SEM. ** denotes *p*=0.007. (G-I) Hemoglobin tetramer HPLC elution chromatograms. Chromatograms represent 2 different donors as well as a technical replicate from donor 2 that was edited independently. (J) Western blot of mock and edited αTM HSPCs at the end of RBC differentiation compared to WT umbilical cord blood-derived RBCs. Western blot image was taken from a single gel that was cropped to place wild-type control next to treated conditions, as indicated by the black line. Loading was standardized by using the same number of cells for input. (K) Ratio of α-globin to dimeric β-globin quantification from western blot. Bars represent mean ± SEM.

To determine whether this strategy was able to restore α-globin production in the context of αTM, we performed HPLC hemoglobin analysis on edited patient-derived HSPCs following RBC differentiation. We found that addition of *tEPOR* to the *α-globin* integration cassette dramatically increased the formation of hemoglobin tetramers, with a substantial increase in HbF, HbA, and HbA2 (Figure 4G-I). Single globin chain HPLC confirmed that α-globin production was indeed elevated in HDR-edited samples (Supplemental Figure 4A-B), likely driving the observed increases in hemoglobin tetramer formation. We next validated these results using western blot for α-globin and β-globin and found that not only do cells edited with *α-globin*+*tEPOR* show a dramatic increase in α-globin compared to unedited and *α-globin* alone-edited patient-derived RBCs, but *α-globin*+*tEPOR*-edited cells also show a decrease in formation of toxic β-globin dimers to nearly WT levels (Figure 4J-K).

### RNA-sequencing demonstrates restoration of oxygen transport and iron ion processes to αTM *HSPCs*

We next used RNA-sequencing to reveal the transcriptional profile of αTM patient-derived HSPCs edited with *α-globin* as well as *α-globin*+*tEPOR* integration vectors in comparison to WT unedited HSPCs. Following editing and *in vitro* RBC differentiation, total RNA was isolated from edited αTM patient cells as well as a healthy WT control. Bulk RNA-sequencing was then performed, followed by quality control and statistical analysis (Figure 5A and Supplemental Figure 5). Total normalized read counts across samples ranged from 17.6-22.0M and mapped to 17,776 unique genes. The high sequence similarity between *HBA1* and *HBA2* precluded clear differentiation; however, reads were exclusively assigned to one gene or another without overlap. For the *EPOR* gene, the alignment was unable to differentiate the endogenous *EPOR* and our transgene *tEPOR*, leading to a combined set of reads from both. In analyzing the data, we found consistent expression across RBC-associated genes in both healthy control and patient-derived cells, which is expected given the efficient erythroid differentiation in edited and unedited samples (Supplementary Figure 6). While the most highly expressed genes by WT RBCs are the adult globins (*HBA1*, *HBA2*, and *HBB*), unedited “mock” αTM RBCs expressed no detectable *HBA1* or *HBA2* (Figure 5B) as expected due to confirmation of four-gene α-globin deletion in this patient. When investigating the genes with the greatest rank change between WT RBCs compared to αTM RBCs, it is unsurprising that the top genes are *HBA1* and *HBA2* (Figure 5C). Despite the modest increase in α-globin production by *α-globin* integration alone according to HPLC, we observed substantial restoration of *HBA1* and *HBA2* expression compared to mock (*HBA1* and *HBA2* rising from undetectable in αTM mock to the 99.9^th^ and 71.7^th^ percentile of expressed genes, respectively) (Figure 5B). Comparing *α-globin*-integrated αTM cells to mock, we found that *HBA1* is the gene that displays the greatest rank change (Figure 5C). Predictably, this restoration of *α-globin* gene expression was further amplified in cells edited with the dual the *α-globin*+*tEPOR* vector, with *HBA1* and *HBA2* expression increasing to the 98.9^th^ and 99.9^th^ percentiles, respectively (Figure 5B). Because this editing strategy places *tEPOR* under the strong, erythroid-specific *HBB* promoter, we also found *EPOR* expression was elevated compared to all other conditions. These elevated expression levels are confirmed by the fact that *HBA1*, *HBA2*, and *EPOR* are among the top four genes with the greatest rank change over unedited αTM cells (Figure 5C).

**Figure 5.**
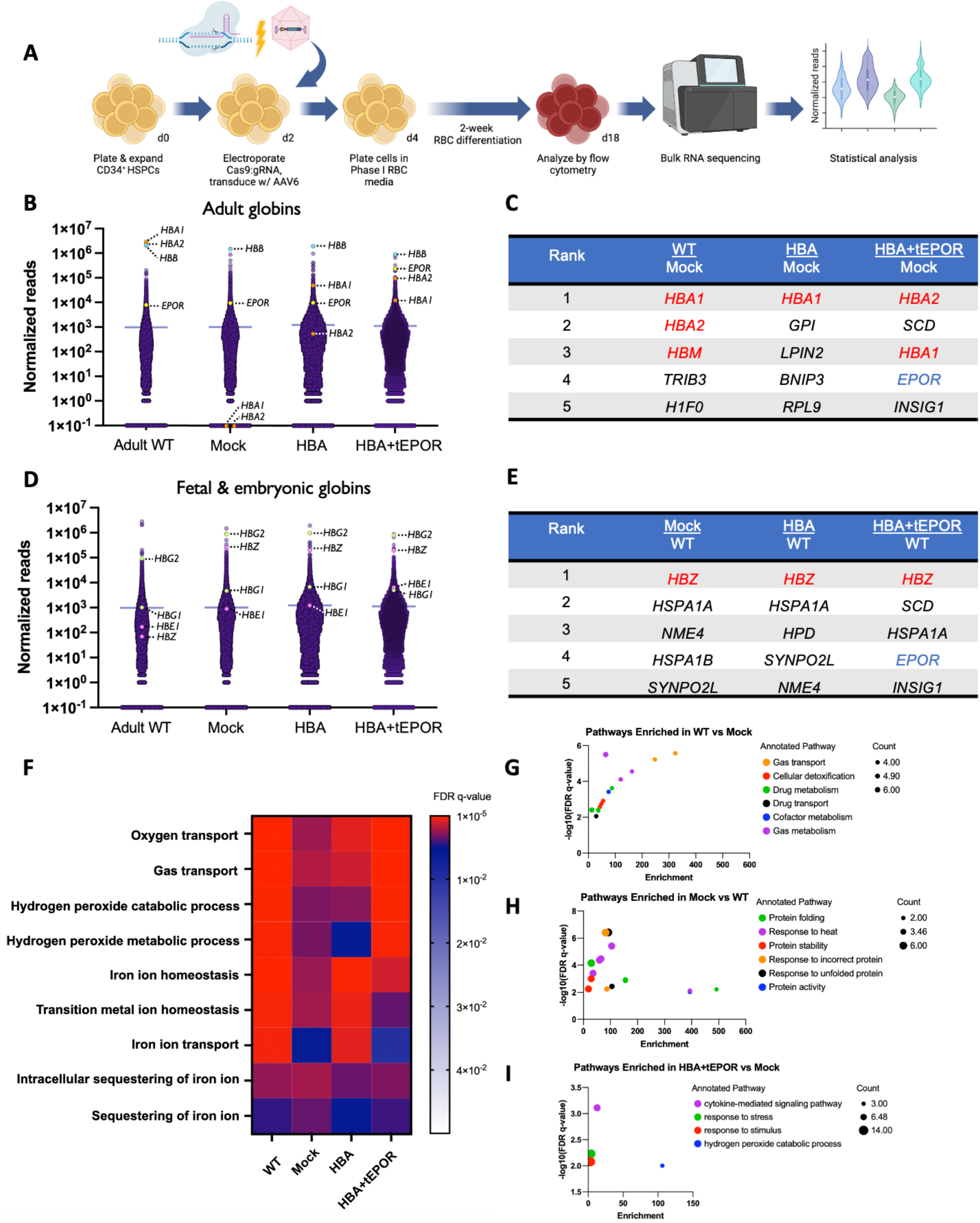
Bulk RNA-sequencing of edited αTM HSPCs demonstrates restoration of α-globin gene expression and oxygen transport and iron ion metabolic processes. (A) Schematic of bulk RNA-sequencing workflow. (B) Normalized reads across all samples displayed with specific genes *HBA1*, *HBA2*, *HBB*, and *EPOR* highlighted. A count of “0.1” was assigned to *HBA1* and *HBA2* for display purposes. (C) Table displaying top 5 genes with the greatest rank change comparing WT and edited αTM cells to mock. (D) Normalized reads across all samples displayed with fetal and embryonic globin genes *HBG1*, *HBG2*, *HBE1*, and *HBZ* highlighted. (E) Table displaying top 5 genes with the greatest rank change comparing mock and edited αTM cells to WT. (F) Heatmap of FDR q-values across 9 commonly enriched GO processes across conditions. Input for GO analysis was the top 0.001% most highly expressed genes within each condition. (G-I) Bubble plots depicting enriched GO processes across treatment comparisons; y-axis displays significance by -log_10_(FDR q-value) and x-axis displays enrichment. Color of bubble pertains to GO process and size of bubble pertains to number of genes represented in the respective GO process. Input for GO analysis was the top 0.001% of genes showing greatest rank change for each comparison (shown in label above each panel).

In addition, we found that *γ-globin* genes (*HBG1* and *HBG2*) were more highly expressed in all αTM samples, regardless of editing, compared to the WT control (Figure 5D). This was expected as the αTM HSPCs were derived from a ∼1-year-old patient, whereas the WT HSPCs were derived from an adult donor. Although ζ-globin is normally expressed only during the first three months of gestation, we found that this gene was highly elevated in all αTM cells (among the eight most highly expressed genes) compared to WT cells (Figure 5D). ζ-globin was also the gene with the greatest rank change in expression in all αTM conditions compared to WT control cells (Figure 5E).

To identify large-scale changes in biological processes, gene ontology (GO) enrichment analysis was performed across the top 0.001% (eighteen) most highly expressed genes across all samples. Compared to unedited mock αTM cells, *α-globin* integration alone led to enrichment of various iron ion-associated pathways to levels equivalent to the WT control (Figure 5F). Further improvements were observed in αTM cells edited with the dual *α-globin*+*tEPOR* integration strategy, showing enrichment of oxygen and gas transport as well as hydrogen peroxide-associated pathways to levels observed in the WT control (Figure 5F). In addition to assessing GO enrichment on the most highly expressed genes within a given sample, we also performed this analysis on the top 0.001% of genes with the greatest rank change across samples. Unsurprisingly, we found that gas transport and gas metabolism were the most significantly enriched pathways when comparing WT to unedited αTM cells (Figure 5G). When performing the opposite comparison, for genes with the greatest rank change in unedited αTM vs. WT cells, we found that the most significantly enriched pathways were for response to incorrect and unfolded protein (Figure 5H), likely representing the cellular response to the formation of toxic β-globin aggregates in the absence of α-globin. Interestingly, when comparing *α-globin*+*tEPOR*-edited αTM cells to WT cells, we saw no significantly enriched GO pathways, indicating that this editing strategy yielded cells that were not appreciably different from normal RBCs. Along with our western blot data (Figure 4I), these findings suggest that while unedited αTM cells have highly elevated β-globin dimers, even a modest increase in α-globin is able to mitigate the unfolded protein response that occurs in the context of αTM pathology. As with WT cells, we also found no significantly upregulated GO pathways when comparing *α-globin*-edited αTM cells to unedited αTM cells (Alpha-thal Supplemental Data). However, when comparing *α-globin*+*tEPOR*-edited cells to unedited αTM cells, we found a number of upregulated GO pathways, the most significantly enriched being associated with cytokine-mediated signaling (likely a result of elevated EPOR signaling) and hydrogen peroxide (H_2_O_2_) catabolism (Figure 5I).

## Discussion

Given the increasing global burden of αTM and tremendous cost of disease management for patients and health care systems, there is an urgent need for curative treatments for this disease. To our knowledge, these results demonstrate for the first time that gene therapy or genome editing may be used to restore normal hemoglobin production to αTM patient-derived RBCs. This approach addresses the two major elements contributing to molecular pathology of the disease in a single genome editing event: successfully increasing α-globin production as well as reducing the formation of toxic β-globin dimers without noticeably disrupting β-globin formation. While the cells in this work were derived from patients with four-gene-deletion αTM, our editing strategy may also be effective in other transfusion-dependent phenotypes of α-thalassemia such as in patients with inactivation of three α-globin genes or hemoglobin H disease^26^.

In addition to these protein-level changes, deeper analysis reveals several interesting findings at a transcriptional level. The enrichment of pathways associated with various oxidative pathways in *α-globin*+*tEPOR*-edited cells suggest an improvement in the stability and function of edited RBCs. Most notably, H_2_O_2_ catabolism and metabolism has been identified as a critical process in preventing cellular injury in β-thalassemic erythrocytes^27^, as the auto-oxidation of unpaired globin chains results in the generation and release of significant amounts of superoxide (O_2_^-^) and H_2_O_2_. These observations suggest that our novel genome editing strategy not only restores *α-globin* gene expression, but also re-establishes essential antioxidant pathways that may be critical in addressing the pathophysiology of α-thalassemia. Moreover, the high expression of ζ-globin found in αTM cells hints at a possible compensatory mechanism for the loss of α-globin in the cell. These findings support the hypothesis that the presence of ζ-globin in patients with α-globin deletions may be protective^4,28^.

In this work, we discovered that pairing *α-globin* transgene with a naturally occurring *EPOR* variant led to the greatest restoration of α-globin production by substantially increasing erythropoietic output of α-globin-expressing αTM patient-derived cells. While we observed significantly higher production of adult hemoglobin tetramers from our *HBA*+*tEPOR* edited conditions, it is clear from our HPLC analysis that fetal-to-adult hemoglobin switching is still underway. Because we are knocking our transgene cassette into the *HBB* locus that is only expressed later in development, we expect the results to be further amplified as hemoglobin switching is completed. This method has the potential to resolve clinical challenges previously encountered in HSCT for the hemoglobinopathies by lowering editing and engraftment frequencies needed to correct RBC disorders. This strategy has the capacity to reduce or eliminate the need for high-morbidity myeloablation currently required to clear the HSC niche, which stand as a major barrier to HSCT safety and efficacy. Moreover, as patients with severe anemia—such as those with αTM —have increased levels of circulating erythropoietin, a *tEPOR*-based genome editing strategy may be especially potent in this context.

Finally, we believe this work stands as a compelling proof-of-concept. However, before this editing strategy is ready for translation into patients, further preclinical studies are needed. These studies include assessing the behavior of edited human αTM HSPCs after transplantation into mice. This may be difficult due to the limited availability of αTM patient HSPCs given the historical prenatal lethality of the disorder as well as the inability of existing mouse models to effectively model human erythropoiesis *in vivo*. Nevertheless, addressing these challenges will be key to paving the way for the clinical application of this promising editing strategy.

In summary, we present results of a comprehensive genome editing approach for the treatment of α-thalassemia that restores α-globin production, reduces the formation of toxic unpaired β-globin aggregates, and restores the transcriptional profile akin to that of a healthy red blood cell. We believe that these findings support development of a definitive *ex vivo* autologous genome editing strategy that may be curative for patients with α-thalassemia.

## Methods

### *In vitro* culture of primary HSPCs

CD34^+^ HSPCs were isolated from umbilical cord blood (provided by Stanford Binns Program) or sourced from Plerixafor- and/or G-CSF-mobilized peripheral blood (AllCells and STEMCELL Technologies). Bone marrow aspirates taken from patients with αTM under protocol no. 16-21157, which was approved by the NHLBI Institutional Review Board. Patients provided informed consent and samples were de-identified after collection. CD34^+^ HSPCs were isolated using a EasySep Human CD34 Positive Selection Kit II according to the manufacturer’s protocol. CD34^+^ HSPCs were cultured at 10^5^ cells/mL in StemSpan SFEM II Medium supplemented with 100ng/mL SCF, 100ng/mL TPO, 100ng/mL FLT3, 100ng/mL IL-6, 20mg/mL streptomycin, and 20U/mL penicillin. Incubator conditions were 37°C, 5% CO_2_, and 5% O_2_.

### Genome editing of HSPCs

Cas9 protein was purchased from Integrated DNA Technologies or Aldevron. Preceding electroporation, ribonucleoproteins (RNPs) were complexed with guide RNAs (gRNAs) at a Cas9:gRNA molar ratio of 1:2.5 at 25°C for 10-20mins. HSPCs were then resuspended in Lonza P3 buffer with RNPs and subsequently electroporated using a Lonza 4D Nucleofector (program DZ-100). Electroporated cells were then plated at 10^5^ cells/mL in HSPC media and AAV6 was added at 5×10^3^ vector genomes/cell based on titers determined by droplet digital PCR (ddPCR). The small molecule AZD-7648 was also added to cells to improve HDR frequencies as previously reported^35^.

### *In vitro* differentiation of HSPCs into erythrocytes

2-3d post-targeting, HSPCs were transitioned into StemSpan SFEM II Medium supplemented with 100U/mL penicillin–streptomycin, 10ng/mL SCF, 1ng/mL IL-3, 3U/mL EPO, 200μg/mL transferrin, 3% antibody serum, 2% human plasma, 10μg/mL insulin, and 3U/mL heparin and maintained at 10^5^ cells/mL. At d7 of culture, cells were transitioned to the above media without IL-3 and maintained at 10^5^ cells/mL. At d11, cells were transitioned to the above media without IL-3 and with transferrin increased to 1mg/mL and maintained at 10^6^ cells/mL.

### Other methods

Detailed explanations of HUDEP-2 cell culture, quantification of genome editing events, AAV6 vector preparation, high-performance liquid chromatography (HPLC), western blot, and RNA-sequencing protocols are described in the Supplemental Methods section.

## Acknowledgements

The authors thank the following funding sources that made this work possible: University of California, San Francisco NIH T32 Research Training in Transplant Surgery Fellowship to S.N.C.; American Society of Gene & Cell Therapy Career Development Award to M.K.C.; NIH NHLBI (no. R01-HL161291) to M.K.C., M.H.P., and T.C.M. and funding from the UCSF Center for Maternal-Fetal Precision Medicine. We thank Emma Canepa and Billie Lianoglou for their assistance with patient sample collection and our patients for their gracious participation in research.

---

**Supplemental Table 1.**
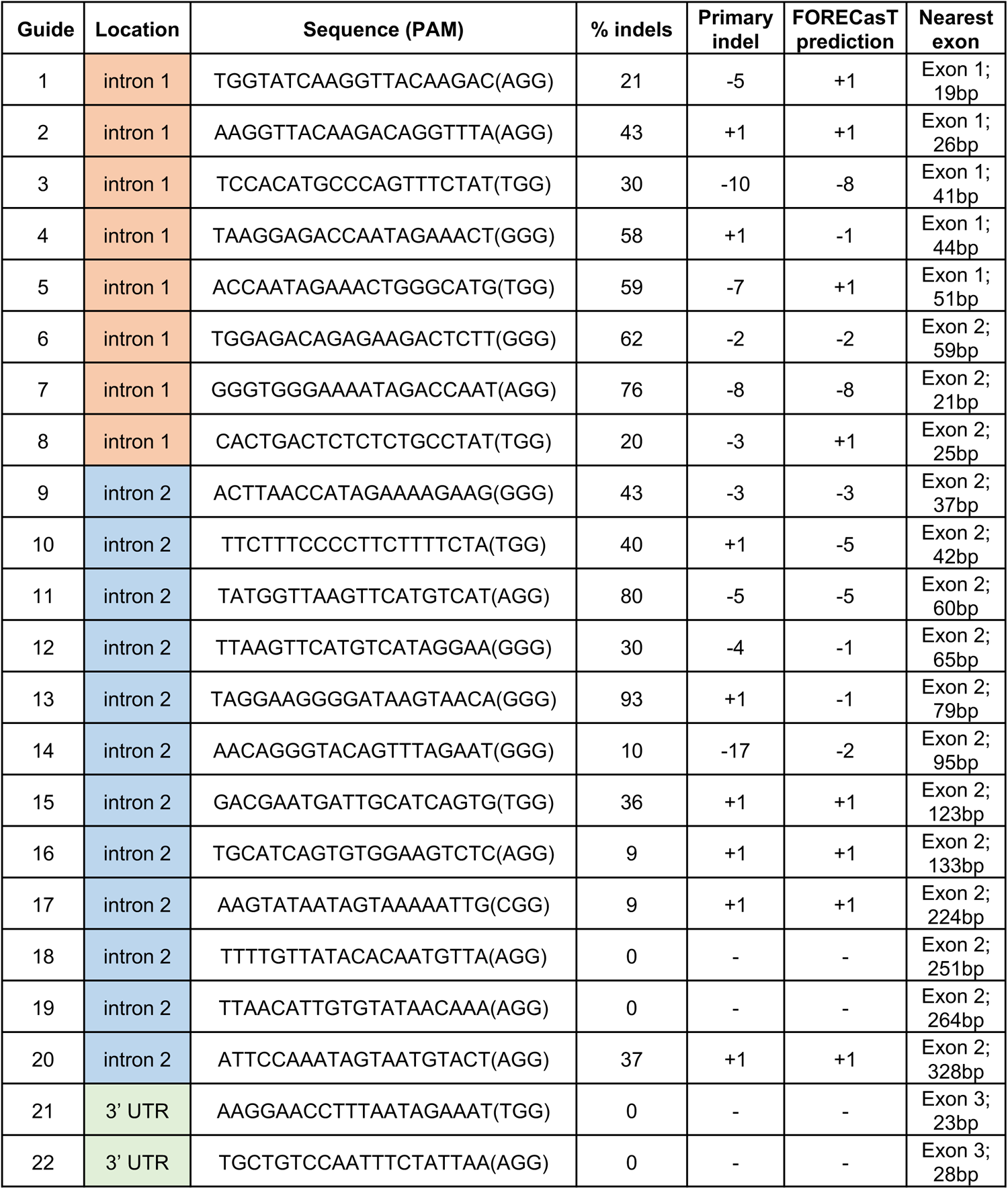

**Supplemental Figure 1.**
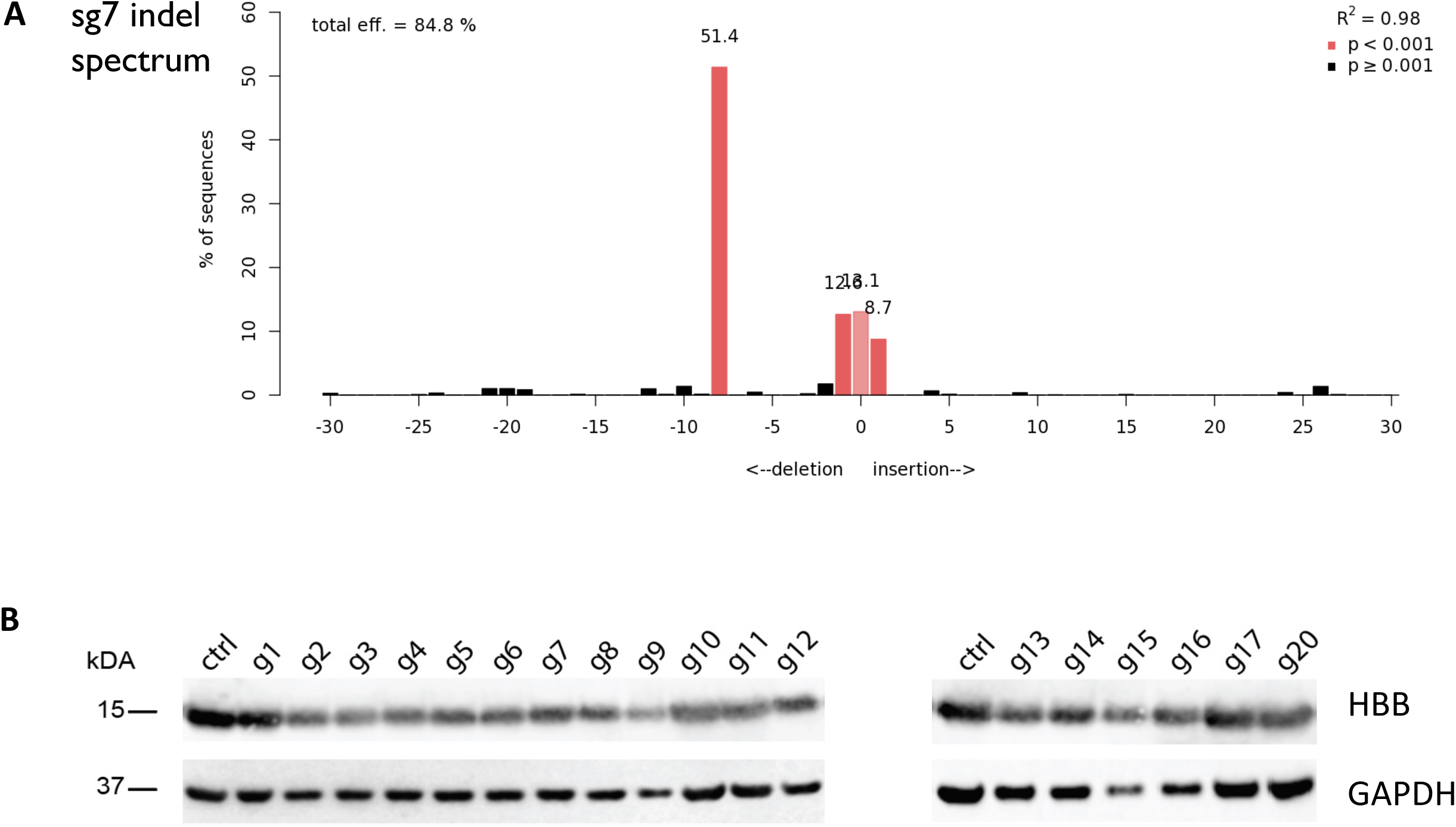

**Supplemental Figure 2.**
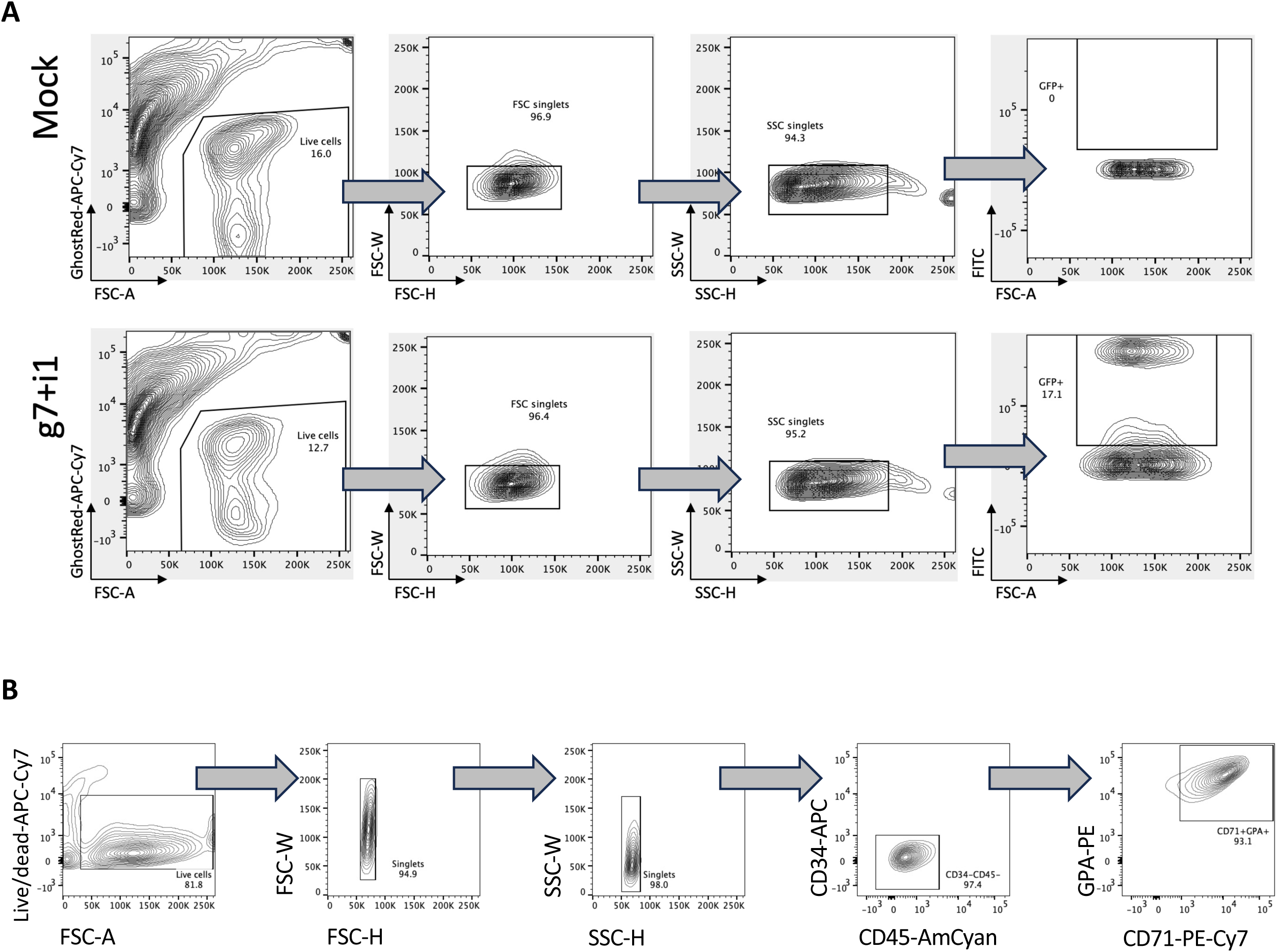

**Supplemental Figure 3.**
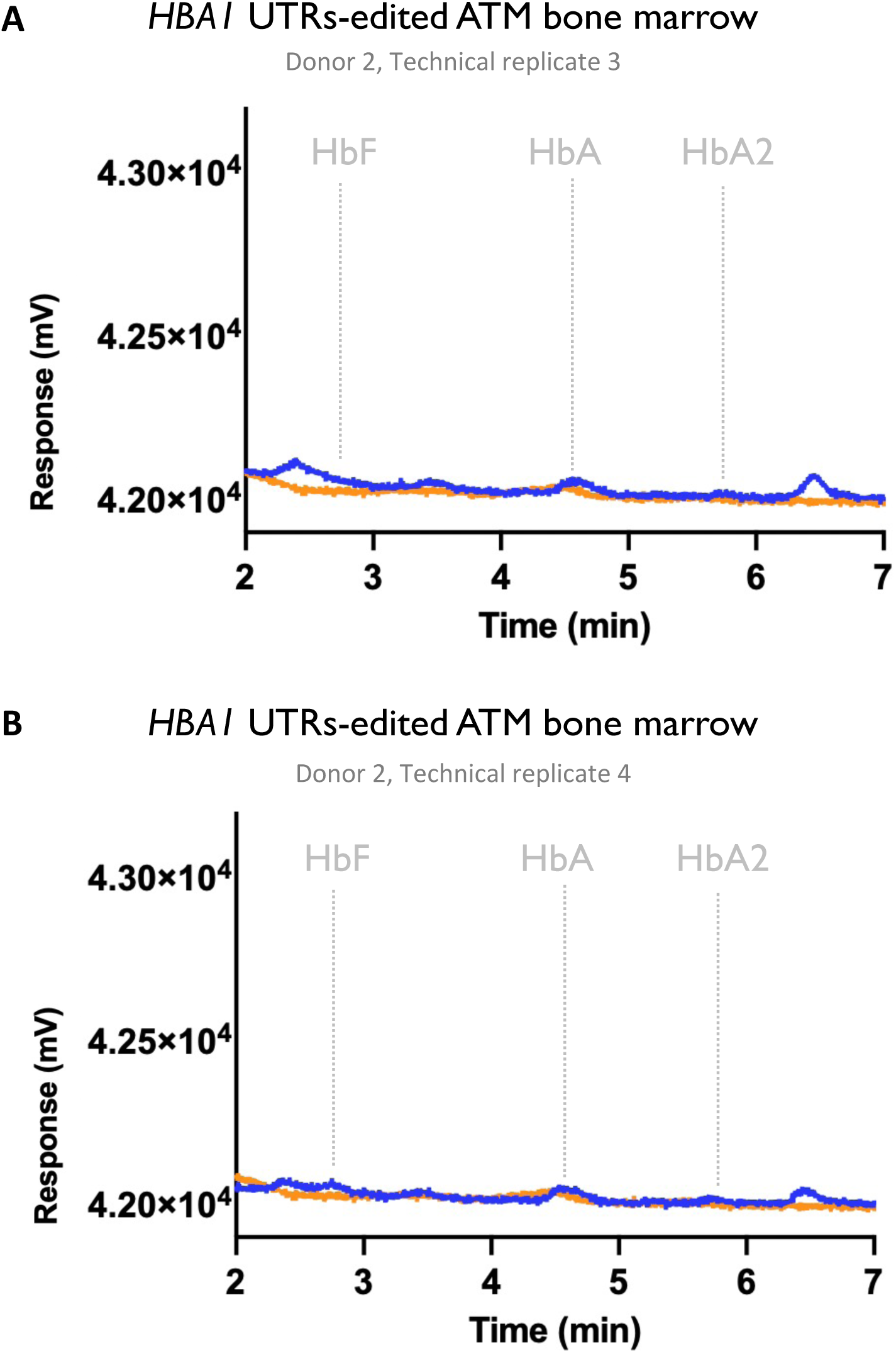

**Supplemental Figure 4.**
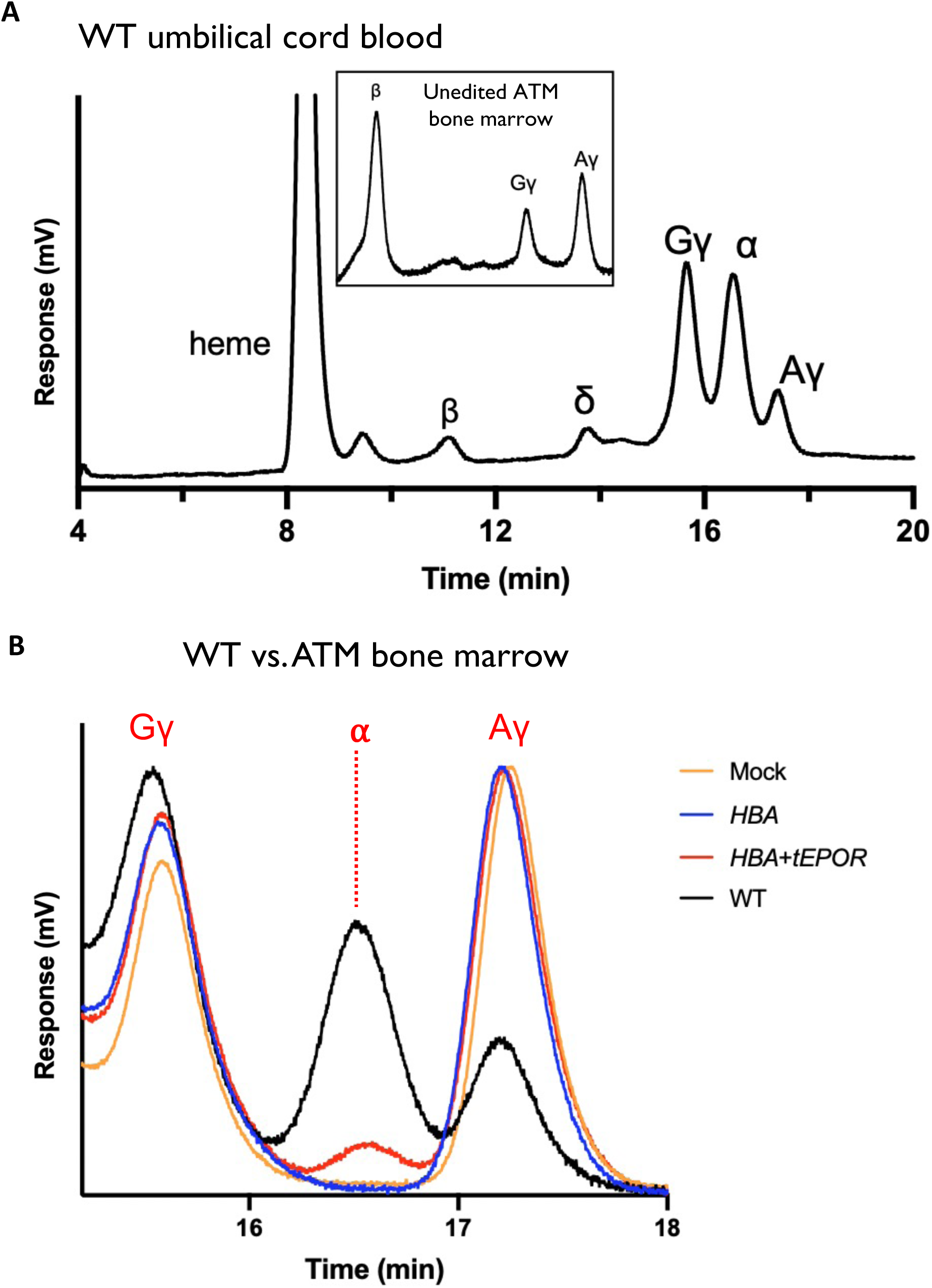

**Supplemental Figure 5.**
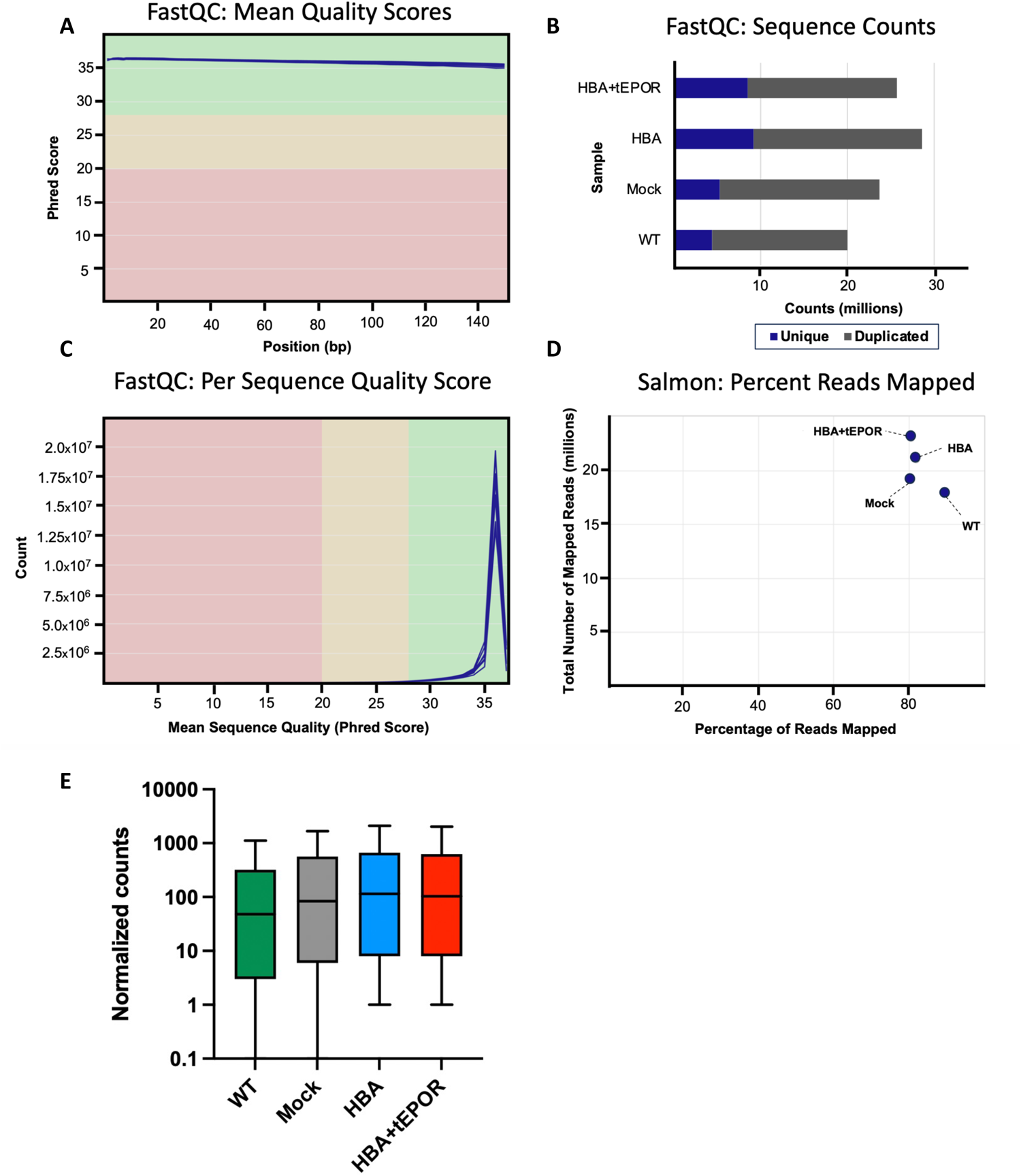

**Supplemental Figure 6.**
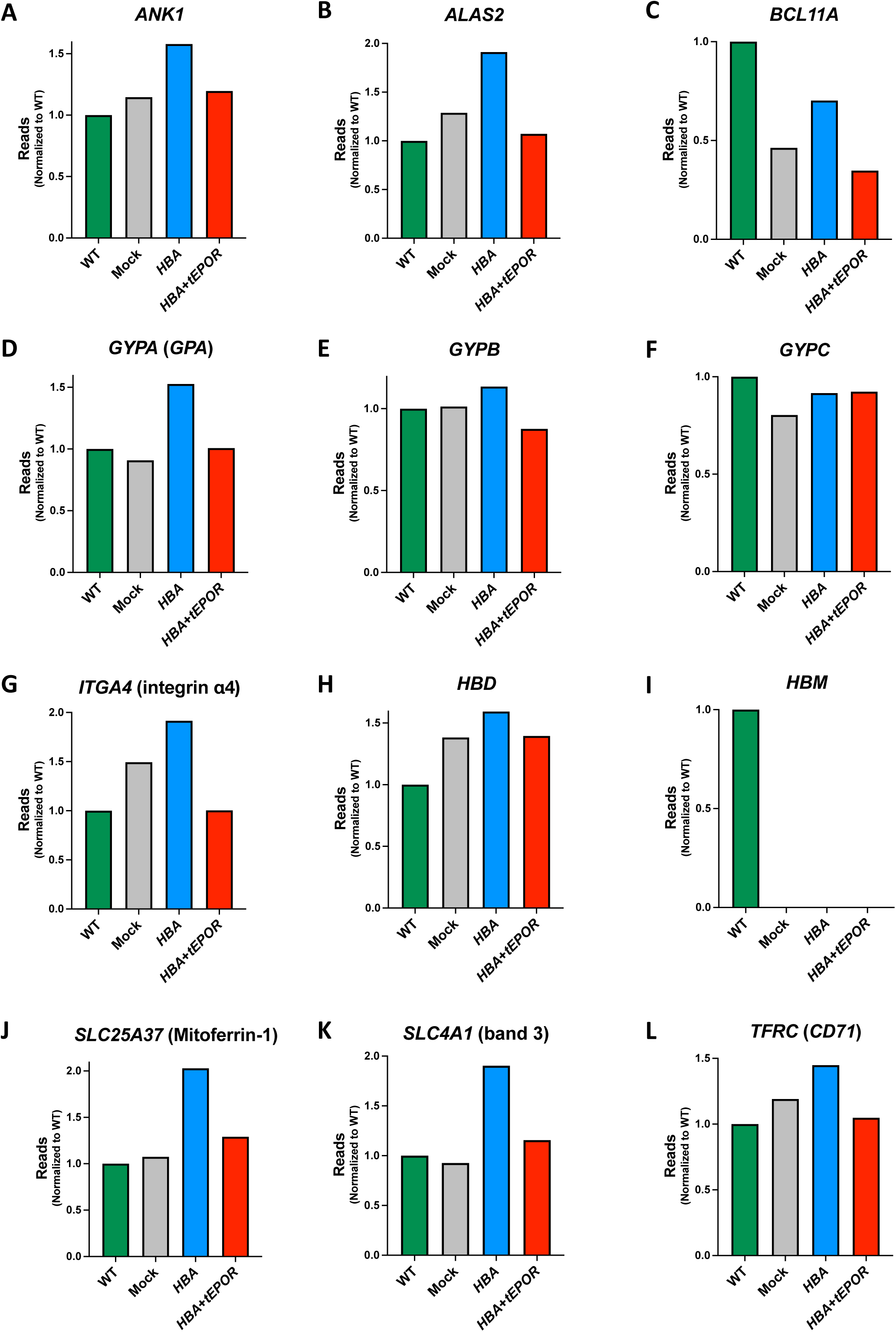

## References

1. Vichinsky E. Complexity of alpha thalassemia: growing health problem with new approaches to screening, diagnosis, and therapy. Ann N Y Acad Sci. 2010;1202:180–187.

2. Piel FB, Weatherall DJ. The α-Thalassemias. New England Journal of Medicine. 2014;371(20):1908–1916.

3. Kreger EM, Singer ST, Witt RG, et al. Favorable outcomes after in utero transfusion in fetuses with alpha thalassemia major: a case series and review of the literature. Prenat Diagn. 2016;36(13):1242–1249.

4. Schwab ME, Lianoglou BR, Gano D, et al. The impact of in utero transfusions on perinatal outcomes in patients with alpha thalassemia major: the UCSF registry. Blood Adv. 2023;7(2):269–279.

5. Schrier SL, Centis F, Verneris M, Ma L, Angelucci E. The role of oxidant injury in the pathophysiology of human thalassemias. Redox Rep. 2003;8(5):241–245.

6. Baird DC, Batten SH, Sparks SK. Alpha- and Beta-thalassemia: Rapid Evidence Review. Am Fam Physician. 2022;105(3):272–280.

7. Fleischhauer K, Locatelli F, Zecca M, et al. Graft rejection after unrelated donor hematopoietic stem cell transplantation for thalassemia is associated with nonpermissive HLA-DPB1 disparity in host-versus-graft direction. Blood. 2006;107(7):2984–2992.

8. Cromer MK, Camarena J, Martin RM, et al. Gene replacement of α-globin with β-globin restores hemoglobin balance in β-thalassemia-derived hematopoietic stem and progenitor cells. Nat Med. 2021;27(4):677–687.

9. Thompson AA, Walters MC, Kwiatkowski J, et al. Gene Therapy in Patients with Transfusion-Dependent β-Thalassemia. N Engl J Med. 2018;378(16):1479–1493.

10. Frangoul H, Altshuler D, Cappellini MD, et al. CRISPR-Cas9 Gene Editing for Sickle Cell Disease and β-Thalassemia. New England Journal of Medicine. 2021;384(3):252–260.

11. Camarena J, Luna SE, Hampton JP, et al. Using human genetics to develop strategies to increase erythropoietic output from genome-edited hematopoietic stem and progenitor cells. Published online August 4, 2023:2023.08.04.552064.

12. Chakraborty N, Bilgrami S, Maness L, et al. Myeloablative chemotherapy with autologous peripheral blood stem cell transplantation for metastatic breast cancer: immunologic consequences affecting clinical outcome. Bone Marrow Transplant. 1999;24(8):837–843.

13. Bhatia S. Long-term health impacts of hematopoietic stem cell transplantation inform recommendations for follow-up. Expert Rev Hematol. 2011;4(4):437–452; quiz 453-454.

14. Inamoto Y, Shah NN, Savani BN, et al. Secondary solid cancer screening following hematopoietic cell transplantation. Bone Marrow Transplant. 2015;50(8):1013–1023.

15. Selvaraj S, Feist WN, Viel S, et al. High-efficiency transgene integration by homology-directed repair in human primary cells using DNA-PKcs inhibition. Nat Biotechnol. Published online August 3, 2023:1–14.

16. Dever DP, Bak RO, Reinisch A, et al. CRISPR/Cas9 β-globin gene targeting in human haematopoietic stem cells. Nature. 2016;539(7629):384–389.

17. Kurita R, Suda N, Sudo K, et al. Establishment of Immortalized Human Erythroid Progenitor Cell Lines Able to Produce Enucleated Red Blood Cells. PLOS ONE. 2013;8(3):e59890.

18. Conant D, Hsiau T, Rossi N, et al. Inference of CRISPR Edits from Sanger Trace Data. CRISPR J. 2022;5(1):123–130.

19. Brinkman EK, Chen T, Amendola M, van Steensel B. Easy quantitative assessment of genome editing by sequence trace decomposition. Nucleic Acids Res. 2014;42(22):e168.

20. Cradick TJ, Qiu P, Lee CM, Fine EJ, Bao G. COSMID: A Web-based Tool for Identifying and Validating CRISPR/Cas Off-target Sites. Mol Ther Nucleic Acids. 2014;3(12):e214.

21. Cromer MK, Majeti KR, Rettig GR, et al. Comparative analysis of CRISPR off-target discovery tools following ex vivo editing of CD34+ hematopoietic stem and progenitor cells. Mol Ther. 2023;31(4):1074–1087.

22. Vakulskas CA, Dever DP, Rettig GR, et al. A high-fidelity Cas9 mutant delivered as a ribonucleoprotein complex enables efficient gene editing in human hematopoietic stem and progenitor cells. Nat Med. 2018;24(8):1216–1224.

23. Dulmovits BM, Appiah-Kubi AO, Papoin J, et al. Pomalidomide reverses γ-globin silencing through the transcriptional reprogramming of adult hematopoietic progenitors. Blood. 2016;127(11):1481–1492.

24. de la Chapelle A, Träskelin AL, Juvonen E. Truncated erythropoietin receptor causes dominantly inherited benign human erythrocytosis. Proc Natl Acad Sci U S A. 1993;90(10):4495–4499.

25. Vichinsky EP. Clinical Manifestations of α-Thalassemia. Cold Spring Harb Perspect Med. 2013;3(5):a011742.

26. Scott MD. H2O2 injury in beta thalassemic erythrocytes: protective role of catalase and the prooxidant effects of GSH. Free Radic Biol Med. 2006;40(7):1264–1272.

27. Gregory GL, Wienert B, Schwab M, et al. Investigating Zeta Globin Gene Expression to Develop a Potential Therapy for Alpha Thalassemia Major. Blood. 2020;136:3–4.

28. Hendel A, Bak RO, Clark JT, et al. Chemically modified guide RNAs enhance CRISPR-Cas genome editing in human primary cells. Nat Biotechnol. 2015;33(9):985–989.

29. Khan IF, Hirata RK, Russell DW. AAV-mediated gene targeting methods for human cells. Nat Protoc. 2011;6(4):482–501.

30. Aurnhammer C, Haase M, Muether N, et al. Universal real-time PCR for the detection and quantification of adeno-associated virus serotype 2-derived inverted terminal repeat sequences. Hum Gene Ther Methods. 2012;23(1):18–28.

31. Eden E, Navon R, Steinfeld I, Lipson D, Yakhini Z. GOrilla: a tool for discovery and visualization of enriched GO terms in ranked gene lists. BMC Bioinformatics. 2009;10(1):48.

32. Allen F, Crepaldi L, Alsinet C, et al. Predicting the mutations generated by repair of Cas9-induced double-strand breaks. Nat Biotechnol. 2019;37(1):64–72.

